# New Insights into Plastocyanin– Cytochrome *b_6_f* Formation: the Role of Plastocyanin Phosphorylation

**DOI:** 10.1101/2025.03.07.641983

**Authors:** Yuval Milrad, Daniel Wegemann, Sebastian Kuhlgert, Martin Scholz, Muhammad Younas, André Vidal-Meireles, Michael Hippler

**Affiliations:** Institute of Plant Biology and Biotechnology, University of Münster, 48143 Münster, Germany; Institute of Plant Science and Resources, Okayama University, Kurashiki, Okayama 710-0046,Japan

## Abstract

In this work we investigated the role of plastocyanin (PC) phosphorylation in photosynthetic electron transfer, focusing on interactions with both cytochrome-*b_6_f* (Cyt*b_6_f*) and Photosystem-I (PSI) in *Chlamydomonas reinhardtii*. While the binding and electron transfer between PC and PSI are well characterized, the interaction between PC and Cyt*f* remains less clear. Using chemical cross-linking combined with mass spectrometry, we identified two potential binding models for PC and Cyt*f*: “Side-on” and “Head-on”. To evaluate electron transfer, we developed an *in vitro* system that allowed oxidized PC, formed via light-driven electron transfer at PSI, to re-oxidize Cyt*f*. Our data shows that a phosphomimetic variant of PC, where phosphorylated PC S49 residue interacts with PetA-K188, displays faster Cyt*f* oxidation, likely optimizing binding and electron transfer between PC and Cyt*f*. Additionally, PC phosphomimetic variants exhibited slower transfer rates than wild type, suggesting that phosphorylation modulates PC’s interaction with PSI. This regulation likely optimizes Cyt*f* oxidation and electron transfer under conditions of low PC availability, such as during high light stress. Overall, PC phosphorylation appears to play a role in fine-tuning electron transfer between PSI, Cyt*b_6_f*, and PC, thereby ensuring efficient photosynthesis in dynamic environmental conditions.

## Introduction

In oxygenic photosynthesis, plastocyanin (PC) acts as a soluble electron carrier which forms a complex with photo-oxidized Photosystem I (PSI) and reduces it. The formation of such complexes between a soluble electron carrier and a major, more stationary, metabolic unit, has been thoroughly studied using various techniques (Holm et al., 1996). Generally, the interface through which the electrons are channeled displays hydrophobic stacking in the proximity of an active metalo-cofactor, surrounded by charged residues that play a role in post-reaction ligand unbinding (Finazzi et al., 2005) and pH-dependency (Kuhlgert et al., 2012). Such is the case of complex formation between PC and PSI in *Viridiplantae* (plants and green algae), in which hydrophobic interactions are formed by two α helices, l and l’, stemming from the loops j and j’ in PsaB and PsaA, respectively (Sommer et al., 2002; Sommer et al., 2004). In the luminal surface of PSI, a positively charged N-terminal domain of its subunit PsaF, enables strong electrostatic interactions (Hippler et al., 1998; Farah et al., 1995; Hippler et al., 1996; Hippler et al., 1999). The formation of a stable PSI–PC complex can be measured by flash photolysis. A laser flash triggers fast intramolecular electron transfer between the bound donor and P700^+^, followed by a slower bimolecular reaction as free PC reduces PSI. The amplitude changes of the fast components relative to donor concentration reflect the binding equilibrium, allowing calculation of the dissociation constant K_D_ (Drepper et al., 1996). The K_D_ for oxidized PC is about six times larger than for reduced PC, raising the midpoint redox potential of bound Pc by 50–60 mV and reducing the driving force for electron transfer. Thus, the structural arrangement of the binding sites allows for differential binding between PSI and reduced or oxidized PC. In this setting, a precise knowledge of the binding scenario is required. Recently, structural validations were obtained via cryo-electron microscopy studies on PSI and PC complexes from *Pisum sativum* (Caspy et al., 2020; Caspy et al., 2021) and *Chlamydomonas reinhardtii* (Naschberger et al., 2022). Interestingly, unlike the extremely conserved luminal plane generated by the surface of PsaA and PsaB, the positions of the lysines (positively charged residues) on the PsaF loop are semi-conserved (Figure 1a), with some residues conserved throughout the entire sub-domain of *Viridiplantae* and some conserved only within each phylum (Naschberger et al., 2022). When aligning their spatial positions (Figure 1b,c), the lysines on residues nr. 12, 16, 19, 23, and 30 are conserved throughout the entire domain, while K24 is present only in plant PsaF loops, and both K20 and K27 are present only in green algae (numbering based on the sequence of the green algae *C. reinhardtii*, ID: P12356, starting from D63 = D1 here). In addition, the conserved acidic patch on PC, formed between residues 42-46 (numbering based on the sequence of *C. reinhardtii*, ID: P18068 starting from D48 = D1 here), is positioned in front of the same conserved residues of PsaF (namely K16 and K23, also termed ‘Northern loop’, Figure 1b,c). Moreover, the non-conserved acidic patch of PC, composed of D54 and the loop formed between residues 58-61, which is shorter in green algae (and some orders of Streptophytes such as Poales and Funariales, for a full alignment, see Supplemental File 1) forms tight interactions with the non-conserved residues of PsaF (i.e. K24/ K20 and K27, also termed ‘Southern loop’, Figure 1b,c), in a perfectly mirrored fashion, emphasizing the importance of these electrostatic bonds.

**Figure 1.**
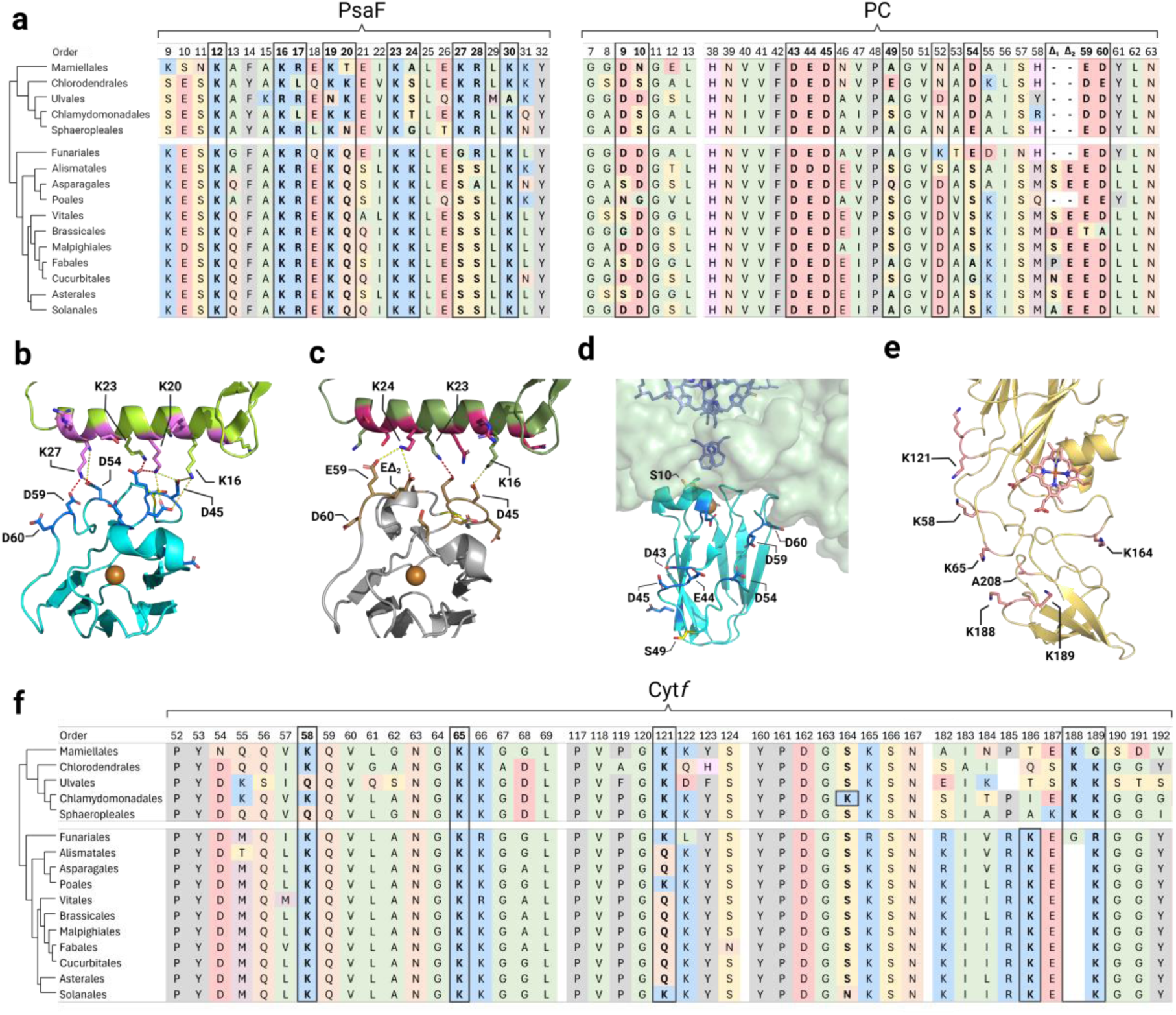
Salt based interaction interphase of algal plastocyanin with photosystem I and cytochrome *b_6_f*. (**a**) Alignment based comparison of relevant loops was conducted for PsaF and PC, using all known sequences (Uniprot.org) of plant (Streptophyta) and green-algae (Chlorophyta) orders. Relevant residues are bolted. (**b-d**) Superposed structure of plastocyanin and photosystem I. Relevant acidic residues (facing the PsaF loop) are presented (blue) as well as two serine residues (S10, S49, yellow), which were shown to be phosphor-regulated. Panels b and c show the formed salt bridges between PC and PSI, comparing algae (**b**, PDB: [7ZQE],[7ZQC]) and plants (**c**, PDB: [6ZOO]) systems. Non-conserved residues of PsaF (green) are highlighted (pink). (**d**) Side view (as viewed from PsaF) of PC from *Chlamydomonas*. *reinhardtii* note that the copper cofactor is situated directly beneath the double tyrosine gateway, leading to the P700 reaction center (purple sticks). (**e**) Structural illustration of Cytochrome *f* (backbone yellow) from *Chlamydomonas reinhardtii* (PDB: [1Q90]), emphasizing its interaction plane and relevant residues (e.g. lysines, salmon). (**f**) Alignment based comparison of relevant loops was conducted for petA (Cyt*f*), using all known sequences of plant (Streptophyta) and green-algae (Chlorophyta) orders. Relevant residues and regions are bolted. The illustration was generated using BioRender.com.

Following its oxidation by PSI, PC is re-reduced by cytochrome *f* (Cyt*f*) of the cytochrome *b_6_f* complex (Cyt*b_6_f*). Here as well, it was shown to form both types of interactions in plants (Ubbink et al., 1998; Sarewicz et al., 2023), where salt bridge formations were shown to hold a major role in stabilizing the complex formation (Soriano et al., 1998; Illerhaus et al., 2000; Gong et al., 2000a). Moreover, specific residues Cyt*f* the hydrophobic patches of both Cyt*f* (Hyun Lee et al., 1995; Gong et al., 2000b) and PC (Illerhaus et al., 2000), were shown to have some essential role in their complex formation. Yet so far, the relevance of the electrostatic interactions could not be confirmed *in vivo* (Soriano et al., 1996), possibly as this reaction is not rate-limiting, in comparison to plastoquinol diffusion and oxidation (Tikhonov, 2024). Still, in many cases, cellular systems use such electrostatic interactions as a regulatory switch by altering the charge of key residues via post translational modifications, such as phosphorylation, acetylation or carbamylation (Balparda et al., 2023). It is quite established, that in plants and algae, active photosynthetic electron transfer activates the kinase STT7 (Depege et al., 2003) via the Qo-site of the Cyt*b_6_f* complex, phosphorylating LCHII proteins and triggering the transition between the so-called State I to State II (Lemeille and Rochaix, 2010). The phospho-proteome of *C. reinhardtii* has been thoroughly studied (Bergner et al., 2015; Ford et al., 2020; Younas et al., 2023; Wang et al., 2014; Werth et al., 2017). Recently, we investigated light-dependent phosphorylation in *C. reinhardtii* and provided evidence that PC becomes phosphorylated *in vivo* and that two phosphorylation sites, namely S10 and S49 were differentially phosphorylated under low (LL) and high light (HL) (Younas et al., 2023). Accordingly, phosphorylation levels of S49 were highly up-regulated under HL, whereas the overall phosphorylation of S10 was down-regulated under similar conditions and in accordance with diminished PC amounts. Interestingly, residue 10 in vascular plants contains an aspartic acid (Figure 1a), which seems to not fit the type of interactions in that region (Figure 1d). In most green algae, the residue is replaced by an uncharged serine (Figure. 1a), which phosphorylation, in a way, mimics the negative charge of the vascular plant type PC (Younas et al., 2023). On the other hand, residue 49 is quite variable although most abundantly hold a serine as well. This, however, must not be over-interpreted, as we so far face data limitations, due to a low number of available algal sequences. Yet, since these two sites (S10 and S49) are located in relevant positions (Figure 1d) they were postulated to have an effect on the kinetics of PC-PSI reduction. To address how the addition of negative charge *via* phosphorylation could affect binding and electron transfer between PC, PSI, and Cyt*f*, we performed functional studies using genetically engineered, recombinantly produced PC. Moreover, we analyzed the binding between PC and Cyt*f* via chemical cross-linking and mass spectrometry, resulting in structural models that are able to describe the possible modes of interactions between these two proteins.

## Results

### The interaction sphere of plastocyanin and cytochrome b₆f

Throughout the photosynthetic clade of *Viridiplantae* (green algae and plants), the extensively positive charged N-terminal domain of PsaF was shown to be crucial for an efficient electron transfer between PC and PSI (Hippler and Nelson, 2021). In addition to PSI, PC can interact with Cyt*b_6_f*, or more specifically the Heme *f*, which is located within Cyt*f*. Again, the two patches of PC seem to have similar objectives in this interaction (Mayneord et al., 2019; Malone et al., 2021). The backbone residues of the soluble region of Cyt*f* are conserved throughout the entire photosynthetic lineage. Here, in resemblance to PsaF, the region is positively charged due to the presence of lysines (Figure 1e). For a comparison’s sake, we aligned the same orders of plants and algae as before (Figure 1f, numbering based on the sequence of *C. reinhardtii*, ID: P23577 starting from Y32 = Y1 here). We observed two lysines, in positions 58 and 65, to be conserved throughout the entire domain (K66 is conserved as well, but it is structurally buried and not facing the interaction plane). In contrast, K121 was featured in almost all available green algae sequences (and some members of plants such as Poales and Funariales) but was replaced by glutamine in plants. The southern loop (between residues 184-192) is, without a doubt, the most variable one. Not only do most of the residues feature no conservation between orders of algae, but the sequence is almost incomparable with other groups of photosynthesizers. Moreover, the length of the loop is largely diverse, with some *Cyanobacteriota* having the longest, uncharged loops. Despite the divergence in sequences, both green algae and plant exhibit two conserved lysines, which are always present on the tip of that loop (although on average, the algal loop is longer in two amino acids), at positions 188 and 189 in green algae, and at positions 186 and 189 in plants (Figure 1f, original numbering is 185 and 187 as in (Ubbink et al., 1998)). Unlike its interactions with PSI, where the copper must be located in a specific position for the electron to pass through the tyrosine gateway, Heme *f* is located near the surface of the luminal face of Cyt*f*. Studies have shown that there are several possible orientations in which PC can form a stable interaction, while placing the copper ligand in the proximity of Heme *f*. In a cyanobacterial mimicking type complex, termed ‘head-on’, the hydrophobic patch of PC interacts directly with the region of the Heme *f* (Dıaz-Moreno et al., 2005). The negative patches have little importance in this case, and it can be explained by repulsion between the negative southern loop and the negativity of the acidic residues of PC. The other, more stable complex termed ‘side-on’, is more well defined and seems to be dominant in plants and algae (also some *Cyanobacteriota*) (Fedorov et al., 2019). In this conformation, salt bridge formation has a crucial impact on the interactions, similarly to the function of the PsaF loop (Illerhaus et al., 2000; Mayneord et al., 2019), where the non-conserved loop of PC (between residues 58-61) interacts with K58 and K65 (also R208, however it is not present in most plants, but replaced with alanine, Figure 1e). The conserved loop of PC (between residues 42-46) interacts with the southern loop of Cyt*f* (between residues 184-192). One more interesting change that was observed specifically in *C. reinhardtii* is a lysine residue located at position 164. This residue is situated on the other side of the negatively charged region and, so far, has not been reported in the literature (Figure 1e).

### Cross-linking analysis grants new insights on Cytb_6_f and plastocyanin interactions

Since we aimed to test the complex formation between Cyt*b_6_f* and PC, we worked in an isolated system from *C. reinhardtii*. Cyt*b_6_f* complexes were isolated using mutants where Cyt*b_6_f* contains a 6xHis-tag at the N’ terminus of *petA*, and PC was recombinantly expressed in *E.coli*. The complexes and proteins were then either reduced, using 5 mM Sodium Ascorbate, or oxidized, using 5 mM ferricyanide. We pre-activated PC proteins using the cross-linker 1-Ethyl-3- [3-dimethylaminopropyl]carbodiimid-Hydrochlorid (EDC) and Sulfo-N-Hydroxysulfosuccinim (NHS) for chemical protein-protein cross-linking, as previously described (Hippler et al., 1989; Hippler et al., 1997; Naschberger et al., 2022). Excess chemicals were then diluted using size exclusion chromatography and additionally washed out via ultra-filtration and centrifugation. Next, the activated PC* was added to the isolated Cyt*b_6_f* mixture. To test whether the cross-linking was effective, we fractionated samples from the mixture via SDS-PAGE and performed both blotting with specific Cyt*f* antibody (Figure 2a, right) and Coomassie staining (Figure 2a, left). The results clearly show that the cross-linking was efficient, as the band of the non-cross-linked PetA subunit (NHS -), which is estimated to be at a size of ∼37 kDa, disappears in the presence of the cross-linker (NHS +) and shifts to ∼50 kDa (as PC is estimated to be at a size of ∼10 kDa). Samples were also digested using trypsin and subjected to a mass-spectrometer analysis (for an elaborated description, see material and methods). Peptide analysis showed several cross-linking events (Figure 1b), mostly peptide PC:59-<u>DD</u>YLNAPGETYSVK, with both PetA: 56-QV<u>K</u>QVLANG<u>K</u>, 59-QVLANG<u>KK</u>, correlated to the cross-linking events of PC:D59/D60 with PetA:K58/K65. In addition, many cross-linking events were also observed with the peptide PetA:111-NILVVGPVPG<u>KK</u>, which indicates on cross-links to PetA:K121, and with the peptide PetA179-IVAITALSE<u>KK</u>, indicating on cross-links to PetA:K188/K189. Surprisingly, the results showed a high abundance of cross-linking events with PetA:157-GQVYPDG<u>KK</u>, suggesting an interaction between PC:D59/D60 from PC and K164 of PetA K66 is conserved as well, but it is structurally buried and not facing the interaction plane therefore present the results as a combination of all identified cross-linked events from 3 independent replicates (for more details, see Supplemental File 2). Interestingly, cross-links with the conserved loop of PC (containing residues D43, E44, and D45), were not detected and so was the peptide containing PC:D54. Indeed, the peptide on which these residues are located has no lysine (which is required for trypsin digestion), resulting in quite a long peptide segment (PC:24-SGETVNFVNNAGFPHNIVF<u>DED</u>AIP<u>S</u>GVNA<u>D</u>AISR). In order to both diminish the size of the peptide and separate the segments containing D43-E44-D45 and D54, we introduced a single point mutation at the residue S49 (which is facing outwards and therefore decreased the chances of miss-folding), altering it to a lysine (S49K). This strategy was proven fruitful (Figure 2c), as cross-linking between this version of PC and Cyt*b_6_f* resulted in many cross-linking events between D54 (PC:50-GVNA<u>D</u>AISR) and all other relevant peptides (except for PetA:K121). In addition, we were also able to detect cross-linked peptides between PC:D45/D54 and PetA:K189/K189, as was expected.

**Figure 2.**
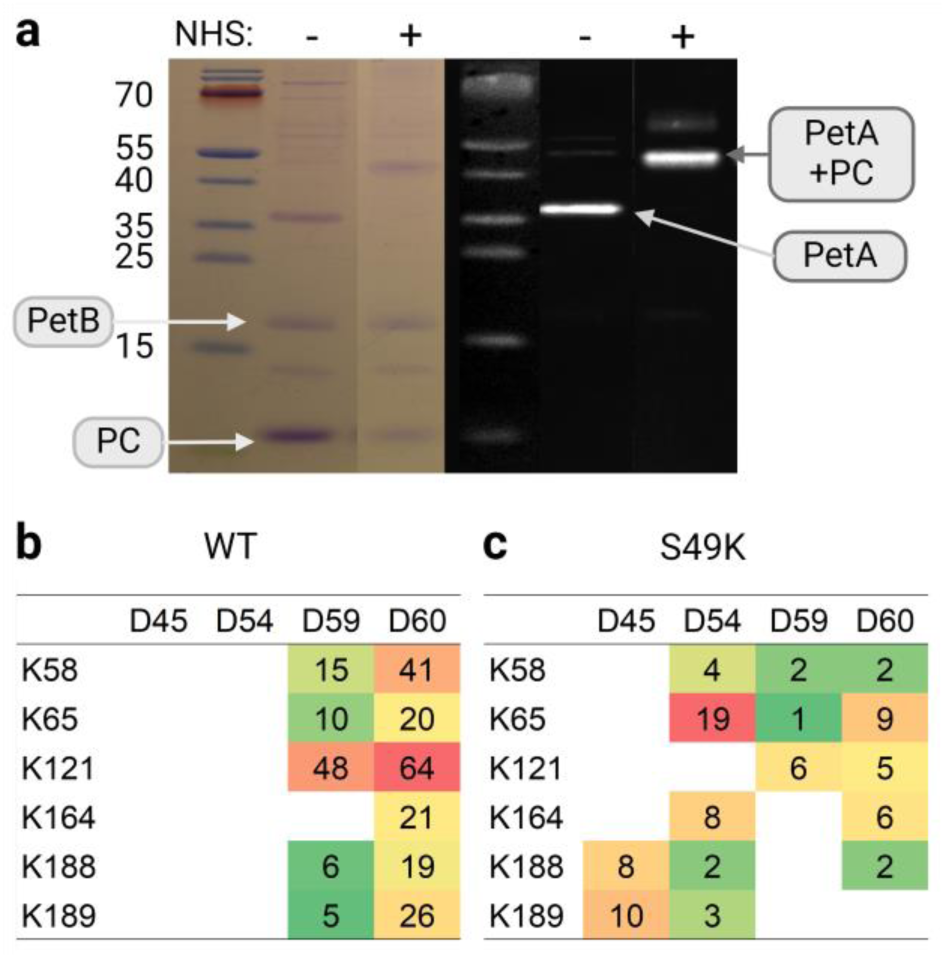
EDC-NHS cross-linking of cytochrome *b_6_f* and PC. Recombinant plastocyanin was activated with EDC and NHS and cross-linked with purified His-tag Cyt*b₆f*. SDS-PAGE fractionation of these samples showed cross-linked (NHS+) and non-cross-linked (NHS-) bands in both Coomassie blue stain and PetA antibodies based Western blots (**a**). The cross-linked samples were digested by trypsin and analysed via mass-spectrometry. Cross-linked peptides revealed an abundant PC:D59/D66 cross-linked to both PetA-K58,K65,K121 and PetA-K164, which is located on the other side of the interaction surface (**b**). Although previous models predicted tight interaction between the southern loop of PC (D43,E44,D45,D54) and PetA (K188,K189), hardly any crosses were detected (WT). We therefore tested recombinant PC mutants with a point mutation at-S49K, generating additional cutting site, and thus enabled a better detection of the relevant peptides (**c**). The illustration was generated using BioRender.com.

### Structural modeling results in shifted interaction conformation

According to the NMR (Ubbink et al., 1998; Lange et al., 2005) and cryo-EM (Sarewicz et al., 2023) structural data, in order to establish a stable complex of Cyt*b_6_f* and PC, the two regions of the acidic patch need to be located in the proximity of the lysines situated on PetA. In plants (Figure 3a, PDB: 2PCF), the PC loop between residues 58-61 is in a tight interaction with PetA Northern loop (distances are specified in Supplemental File 3, structural alignments are shown in Supplemental File 4). The conserved PC loop (between residues 42-46) interacts with the Southern PetA loop (residues K186, K189). In addition to that, PC:D52 (in plants) interacts with both PetA:K189 and R208, which as mentioned, can only be found in some plant’s lineages (and interacts with D43 as well). This complex formation led to a core/core distance (between the copper atom and the iron center of Heme *f*) of 11.9 Ȧ. Alignments of algal type PC and PetA, based on the location of the copper and backbone (Figure 3b, PDB: 1Q90, 7ZQE) results in several issues. For one, the shortened non-conserved loop of PC (residues 58-61) cannot access the positive region and form bonds with PetA:K58, even if PC:D59 is in the proximity of PetA:K65. Moreover, R208 is replaced by alanine, resulting in diminished interactions for PC:D54 (which resembles in a way the position of D52 in plants). Bonds between the Southern loop of PetA and PC are quite similar, however a simple alignment results in a clash between PetA:K188 and PC:E44. In order to tackle these issues, we attempted to improve the model by using a modelling server, ClusPro.2 (Kozakov et al., 2017; Desta et al., 2020). This server ensures an energetically stable alignment of the two given proteins, based on bond formations between them (in all forms). To specify optimal complex formations, we restricted some of the interactions to a hypothetical distance between relevant residues (Supplemental File 3, (Xia et al., 2016)). Since we based our model on plant interactions, we began by restricting interactions between PetA:K58,K65 and PC:D59,D60. To mimic the natural salt bridge formation of this complex, we determined the restricted distances to be between 1-5 Ȧ, with 50% chance of required bond formations (see Supplemental Files 4, nms_01). This however resulted in a distortion of the two acidic residues to be locked on PetA:K65, as their reach was not good enough to form bonds with PetA:K58. Unsurprisingly, the Southern loop of PetA was prominent to the complex formation, as PetA:K189 interacts with PC:D54 and PetA:K188 interacts with PC:E44 (see Supplemental File 4, nms_01). However, the distortion led to an increased distance between the cores to 13.4 Ȧ. Indeed, even in case that the orientation of the Cu center along the Heme plane generate no effect the possible electron transmission, the increased spatial distance will result in it being less effective (Pietras et al., 2014), which seem unlikely due to the evolutionary needs of the organisms and deem the model less viable. To entangle this issue, we reevaluated the restrictions posed by the software, in accordance with our mass-spectrometry observations (for a full description, see Supplemental File 3, structural alignments are shown in Supplemental File 4, nms_02). As mentioned before, algal type Cyt*b_6_f* has a lysine at position 121, which resulted in many cross-linking events with PC:D59 and D60 (Figure. 2). In addition, we took into consideration the possibility of bond formations between PC:D54 and PetA:K65 and interactions via the Southern loop of PetA (K188 and K189) with the conserved region of PC (D43,E44 and D45). This model (Figure 3c, see also Supplemental Figure 1) results in a core distance of 10.9 Ȧ, as PC twists to the other side, mainly due to the bond formations with PetA:K121(see Supplemental Figure 1b), positioning PetA:K65 in-between PC:D54 and E44 and place PetA:K188 in the proximity of S49 (see Supplemental Figure 1 c,d), which was, as mentioned, shown to be phosphoregulated. According to this model, the interaction which is established resemble a “side-on” model and the formed complex is likely to be very stable. However, since we also observed a lysine in position 164 of PetA (in Chlamydomonas), which is easily cross-linked to PC:D60/D59 as observed in our results (Figure 2), we opted to decipher another type of interaction, this time restraining the software to take this interaction under consideration (structural alignments are shown in Supplemental File 4, nmh_07). The resulting model showed very interesting features (Figure 3d, see also Supplemental Figure 1). The distance between cores was also relatively short, at 10.9 Ȧ, and in general this model resembles much a “Head-on” type complex. The total bond strength of this model is weak, as in addition to PetA:K164-PC:D60 (see Supplemental Figure 1f) bond we observed only a PetA:K65-PC:D9 and PetA:K121-PC:S10 bonds (see Supplemental Figure 1g). Forming such bonds could be favorable in the case where S10 holds a phosphate group, as was previously discussed.

**Figure 3.**
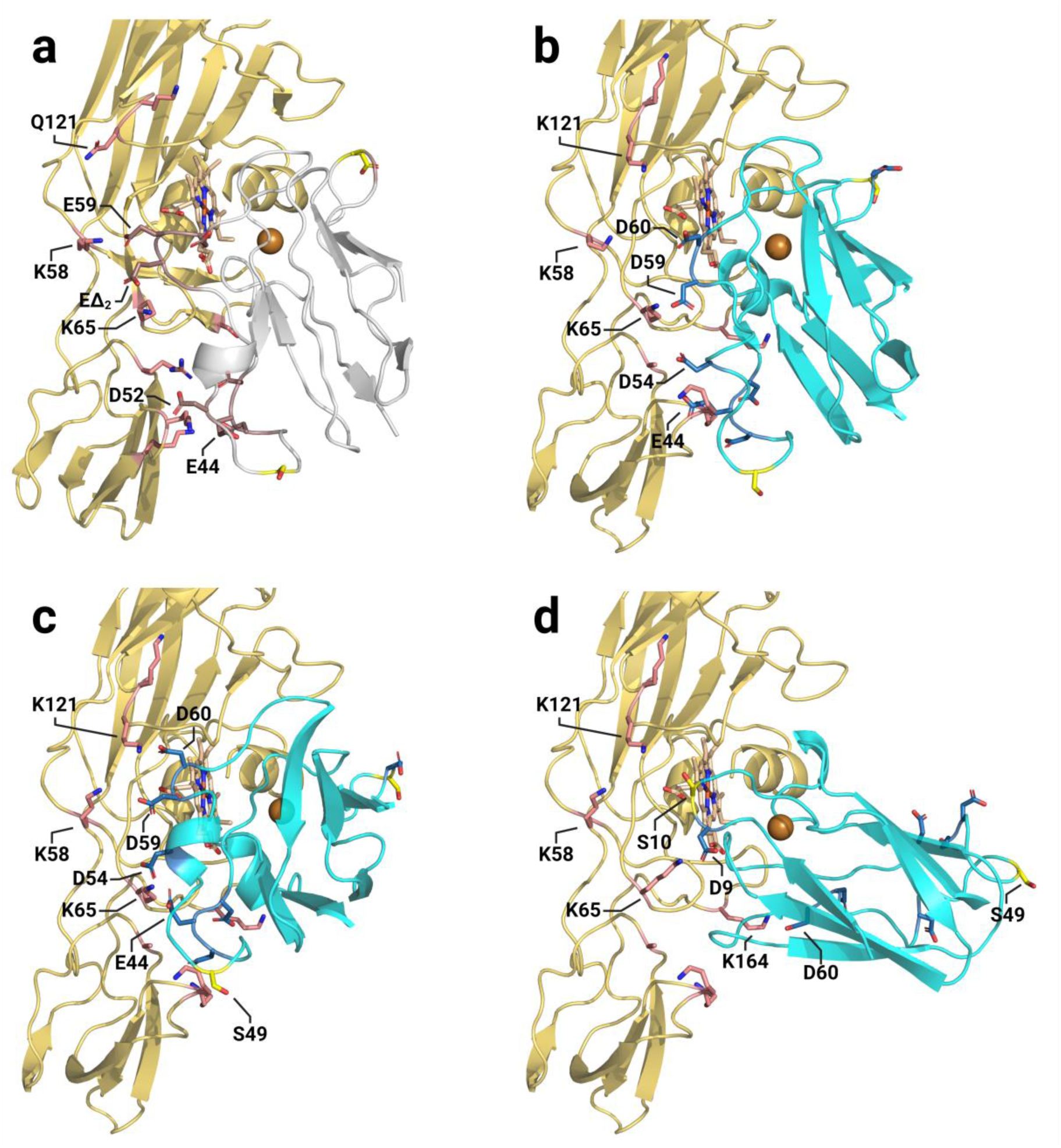
Structural modelling of Cyt*f* and PC in complex. By superposing algal plastocyanin [PDB 7ZQC] to the NMR based plant interaction model [PDB 2PCF] (presented in panel **a**), we tested the distances between the negatively-charged PC resides (blue) and positively-charged Cyt*f* residues (salmon). In accordance with this model, we measured the distance between the metal cores of the proteins (PC-Cu/PetA-heme-Fe) and determined them to be 11.9 Ȧ (**b**). The second model (**c**) takes into consideration the fact that algal PetA has an additional Lys at position 121, and that it was cross-linked to PC:D59/D60. Additional interactions are observed between PC:D54 and PetA-K58/K65 (as was shown in the mutated S49K PC), PC:D43/E44 and PetA-K188/K65. In addition, PC:S49 seems to be in the proximity of PetA-K188 and given a phosphorylation form might increase the stability of the complex formation. Core Cu-Fe distances are predicted to be 10.9 Ȧ. The third model (**d**) takes into consideration a possible interaction between PC:D60 and PetA-K164. This residue cannot be found in plants or *Cyanobacteriota*, but only in green algae. Since we observed such cross-linked peptides, we aligned the residues and postulated that the chances for this orientation to dominate the interaction mode increases under conditions in which PC:S10 is phosphorylated, and PC:S49 remains un-phosphorylated. Core Cu-Fe distances are predicted to be 10.9 Ȧ, which should increase the electron transfer rate. In addition, other negative residues of plastocyanin are not in interaction, which should decrease the complexes strength. All models are available on Supplemental File 4. The illustration was generated using BioRender.com.

### Kinetics assessments of the role of phospho-regulation on plastocyanin

In our structural modelling approach, we observed two possible models that could result in a stark complex between Cyt*b_6_f* and PC, which we termed as ‘Side-on type’ (Figure 3c) and ‘Head-on type’ (Figure 3d) complexes. We also observed that each of these complexes rely on different bonds between the acidic patches of PC and lysine residues situated on PetA and that phosphorylation of PC:S10 or S49 residues could have an impact on their formation. In order to test this hypothesis, we used site-directed mutagenesis and expressed the following recombinant PC variants: PC:S10A, PC:S10E and PC:S49D. These PC variants were compared to WT in P700^+^ re-reduction experiments (Figure 4). To do that, we mixed isolated PSI complexes from *C. reinhardtii* with increasing amounts of PC (0.3-6.0 µM), and measured changes in absorbance (705-740 nm, (Kuhlgert et al., 2012)) following a laser single-turnover flash (Laser type supplied with a red dye, Sigma), using a ‘Joliot type Spectrophotometer’ (JTS -150, Biologics). Detection resolution was set to have 100 measurements during 5 seconds post laser flash, with an initial delay of 700 µsec, since the laser flashes tend to ‘blind’ the detectors in this set-up. Our data show that with increasing concentrations of PC, the rate of P700^+^ re-reduction is increasing (Figure 4a). We then plotted the graphs and fitted them using a double exponential kinetics (Origin.Pro, ExpDec2). K₂ values were calculated according to (Drepper et al., 1996), and the results were plotted in regards to PC concentration (Figure 4b). To compare the differences between PC types, we averaged K₂ values, between 1.0-6.0 µM (in low concentration there seems to be some additional effect, as K₂ is not constant), and plotted the result as box plots (Figure 4c). The results show that all proteins can reduce PSI. Moreover, S10A show no effect on PC-PSI electron transfer rates, while S10E and S49D have a slightly decreased rate.

**Figure 4.**
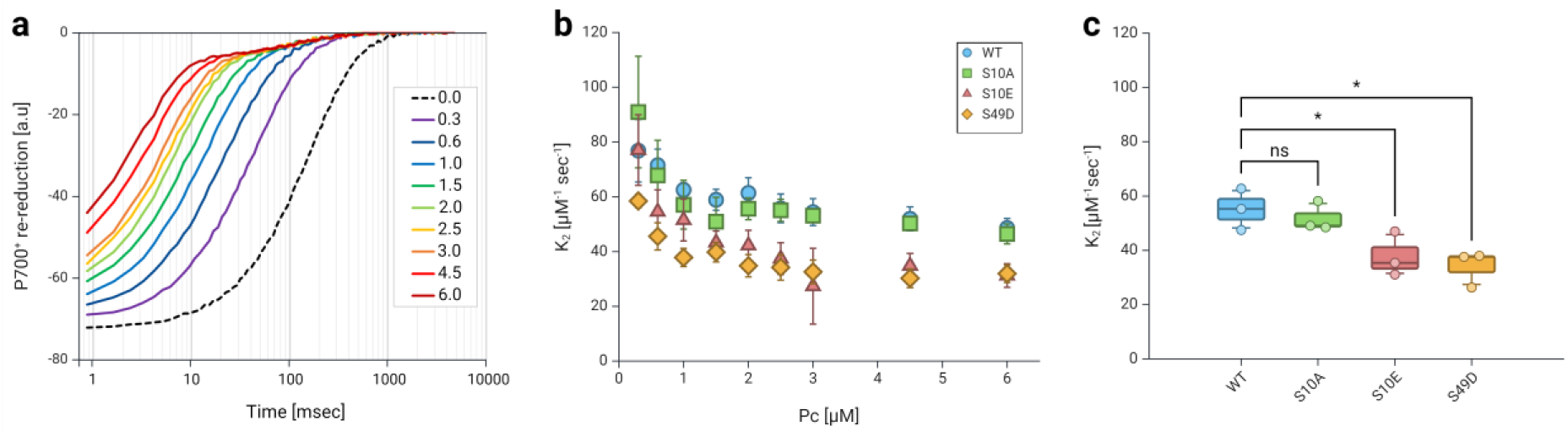
P700^+^ re-reduction curves following laser flash show decreased interaction of phosphorylation mimicked PC. Purified PSI complexes were tested in JTS. (**a**) Following a laser flash, maximal oxidation of P700^+^ centers were determined, followed by a double exponential re-reduction phase. The complexes were mixed with increasing PC concentration and re-reduction rate was increased accordingly. The kinetics featured two phases of reduction, in which the initial reduction, below measurement resolution was detected. To test the effects of phosphorylated plastocyanin variants on the kinetics of P700^+^ re-reduction, the experiment was conducted in the presence of recombinant PC variants (**b**), in which S10 or S49 were replaced by either Ser (WT), Ala (S10A), Glu (S10E), or Asp (S49D). K_2_ values were calculated, using OriginLab, for each concentration of PC. (**c**) Box plots show the averaged K_2_ values of 1-6 µM plastocyanin (3 repetitions). Here, S10A showed no significant change, while both phosphor-mimicking mutations showed a slight decrease. Statistical analysis was conducted using a One-way ANOVA with Dunnett multiple comparisons test. The illustration and statistical analysis were generated using BioRender.com.

### In vitro electron flow shows ferredoxin dependent cytochrome b_6_f activity

Unlike PC-Cyt*b_6_f* complex binding and electron transfer reactions, the kinetics and mode of interaction between PC and PSI are thoroughly researched and very solid. One reason for this difference is the simplicity of measuring PSI activity. PSI oxidation can be triggered by light, as re-reduction via PC can be readily measured. In contrast, Cyt*b_6_f* is not directly oxidized by light and therefore kinetic studies had to come up with more creative measurement set-ups, i.e. measured using fast mixing stopped-flow instruments, taking advantage of differences in its absorbance at 554 nm (with 546 nm and 574 nm usually serve as a reference baseline). In order to tackle this, we took advantage of the properties of PSI as a PC oxidizer. To establish a direct measurement of Cyt*f* oxidation, we mixed 2 µM of isolated Cyt*b_6_f* complexes and 331 nM PSI solubilized in 7.5 mM KCl, 2.5 mM MgCl_2_ and 10mM ascorbate, pH 7.0 (MOPS). In this assay, methyl-viologen, that has an absorbance between 500-600 nm, was replaced with recombinant FDX, based on recombinant PETF from *C. reinhardtii* (Marco et al., 2019), which was also postulated to have the function of Cyt*b_6_f* reducer, as illustrated in (Figure 5a). Initially, we tested the effect of FDX on the kinetics of PSI re-reduction (Figure 5b). The results show that the addition of FDX does not influence the kinetics of P700⁺ re-reduction. Assessments of K₂ did not yield significant differences between the values due to the altered protocol (Figure 5c, FDX+ Cyt-vs. FDX-Cyt-). Moreover, in our experimental set up, we saw a slight decrease on K₂ due to the addition of Cyt*b₆f*, in the presence of WT PC (Figure 5c, FDX+ Cyt-vs. FDX+ Cyt+). We therefore continued to evaluate the effect on the oxidation of Cyt*f*, by measuring its absorbance (Figure 5d, for full data set, see Supplemental File 5), using JTS (JTS-150, Biologics). We added 1 µM PC to the mixture, which was kept in darkness (grey background), before we illuminated it for two seconds (white background), followed by 15 seconds of darkness (4 technical and 3 biological repetitions for each test). In the light phase we observed a stark decrease in Cyt*f* absorbance, which lasted ∼30 ms, followed by a second phase in which it continued to decrease more moderately until a steady state was achieved. As light was turned off, the trace showed an abrupt drop, followed by a steady, exponential increase, back to the original reduced state. We then added increasing concentrations of FDX (0.5-5 µM) and resumed the measurements.. The results show no significant effect of the FDX concentration on the initial stark oxidation at light onset. In contrast, the rate of the second phase of Cyt*f* oxidation seems to be diminished, meaning a more net-reduced Cyt*f*, possibly by FDX. This could indicate on a circular electron flow, as postulated. However, one should take into consideration the relatively low redox potential of FDX, that could definitely reduce the Heme *f* centers directly. Notably, at the onset of darkness, the drop showed no significant changes, nor did the rate in which Cyt*f* achieved re-reduction. We then added increasing amounts of PC to the mixture (up to 3.0 µM) and observed a mirrored effect on the traces (Figure 5e). The rate in which Cyt*f* is oxidized increased, in accordance with the apparent concentration of oxidized PC (which itself is oxidized by PSI). The rates of Cyt*f* dark re-reduction were also affected, as the half time of the reaction increased in accordance with PC concentration, indicating on a lasting effect on Cyt*f* oxidation.

**Figure 5.**
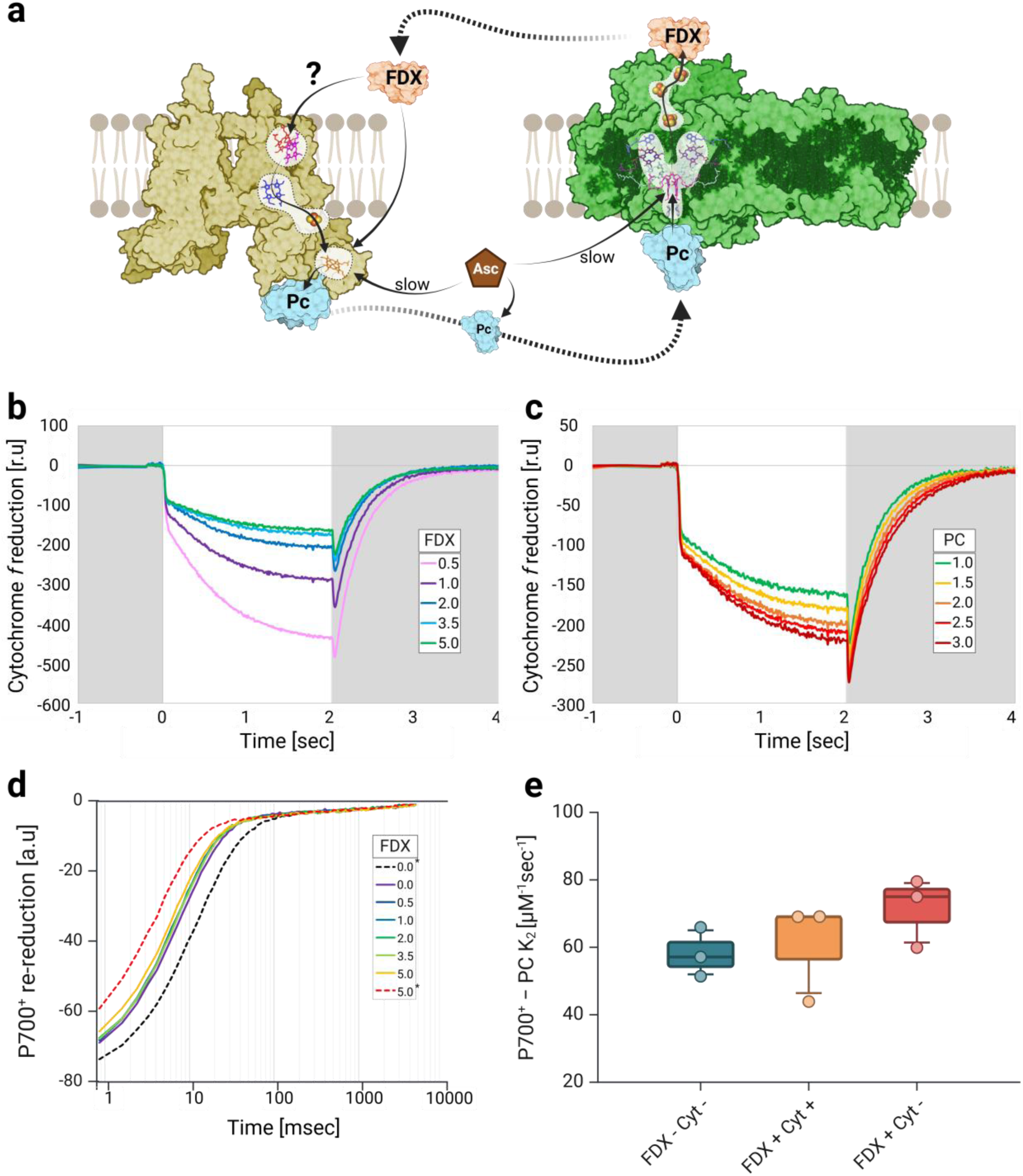
Cyt*f* oxidation kinetics. (**a**) Schematic overview of the cyclic electron chain between Cyt*b_6_f* and PSI via PC and FDX in our experimental set-up. Kinetic measurements were conducted using JTS on isolated Cyt*b_6_f* (2µM) and PSI (300 nM) complexes, in the presence of: 1mM ascorbate and 2.5mM MgCl₂ at pH 7.0. (**b**) Post laser flash P700 measurements were conducted, with increasing concentrations of FDX, in the presence of 1 µM PC (WT, solid lines). Before and after FDX additions, PC was added to illustrate the range of expected changes (dashed lines), with PC additions from 0.6-1.0 µM (0.0*, black) and from 1.0-1.5 µM (5.0*, red). (**c**) K₂ values of P700⁺ re-reduction rates were calculated using OriginLab, were and presented in as box plots, in the presence/absence of FDX (FDX+/-), where FDX-was supplemented with MV (blue). The effect of Cyt*b₆f* complexes was also tested (Cyt+, orange). Results show no significant differences between these treatments. Cyt*f* oxidation kinetics were also tested, as complexes were exposed to 2 seconds of illumination (white background). (**d**) Increasing FDX concentrations, showed decreased oxidation kinetics, possibly due to a faster re-reduction of cytochrome complexes. (**e**) Increasing PC concentrations (in the presence of 5 µM FDX) featured an increase of the second light-phase net oxidation rate and a rapid dark re-reduction (gray background). The illustration was generated using BioRender.com.

Our next step was to test whether differences in Cyt*f* oxidation as a result of the single PC mutations using PC: S10A, S10E and S49D could be observed. To do so, we first assayed P700^+^ re-reduction. Here, we observed a significant decrease in P700^+^ re-reduction rates for both S10E and S49D, suggesting that in the presence of Cyt*b_6_f*, re-reduction of P700^+^ via S10E and S49D is still slowed down, as observed (Figure 6a, orange). Notably, the re-reduction of P700⁺ was diminished furtherly for these PC variants, compared to the decrease of the K₂ values in the absence of Cyt*b₆f* (Figure 6a, blue, see also Figure 4c). These observations hint on a stronger interaction between these PC variants to Cyt*f*, diminishing the apparent PC concentration that could reduce PSI. We then examined the traces of the Cyt*f* oxidation assay (Figure 6b). As before, we fitted the curves to a double exponential regression (Origin.Pro, ExpDec2, for further data see Supplemental File 5, sheet: Fits CytF-light). We observed no significant differences in the total amplitude of these traces between PC variants (nor the values of the amplitude constants, A1 and A2 of the double exponential fit we conducted). We therefore focused our views on the trends of the curves, rather than their amplitudes. To do so, we extrapolated the curves in relations to their steady state oxidation values (following around 2 seconds of light exposure). As no differences were observed in these trends for the increasing PC concentrations (the amplitudes are different, but the rates and trends remain the same per variant), we averaged the relative traces for each variant (between 1-3 µM of PC, with 5 µM FDX), and presented the results in (Figure 6b). The results clearly show a more moderate initial fast drop at the onset of illumination for both S10A and S10E. However, they did not result in any differences in-between these two variants (Figure 6b). Moreover, the second phase seem more prominent in both of these PC variants, compared to the WT, but resulted in no significant differences (Figure 6c). Finally, the dark re-reduction of Cyt*f* was also indistinguishable in rate for these two variants (Figure 6d). In contrast, the addition of PC:S49D showed completely different phenotype, as it lacked the initial drop and phase separation at the onset of illumination (Figure 6b). Instead, the signal decreased rapidly without showing any breaks until it reached steady state with an average t_½_ of ∼120 ms (compared to ∼830 ms of the WT, see Supplemental File 5, sheet: Fits CytF-light). In addition, the dark re-reduction of Cyt*f* was not slowed down by increasing PC:S49D concentrations, as was observed for WT PC, PC:S10A and PC:S10E (Figure 6e, see also Supplemental File 5, sheet: Fits CytF-dark).

**Figure 6.**
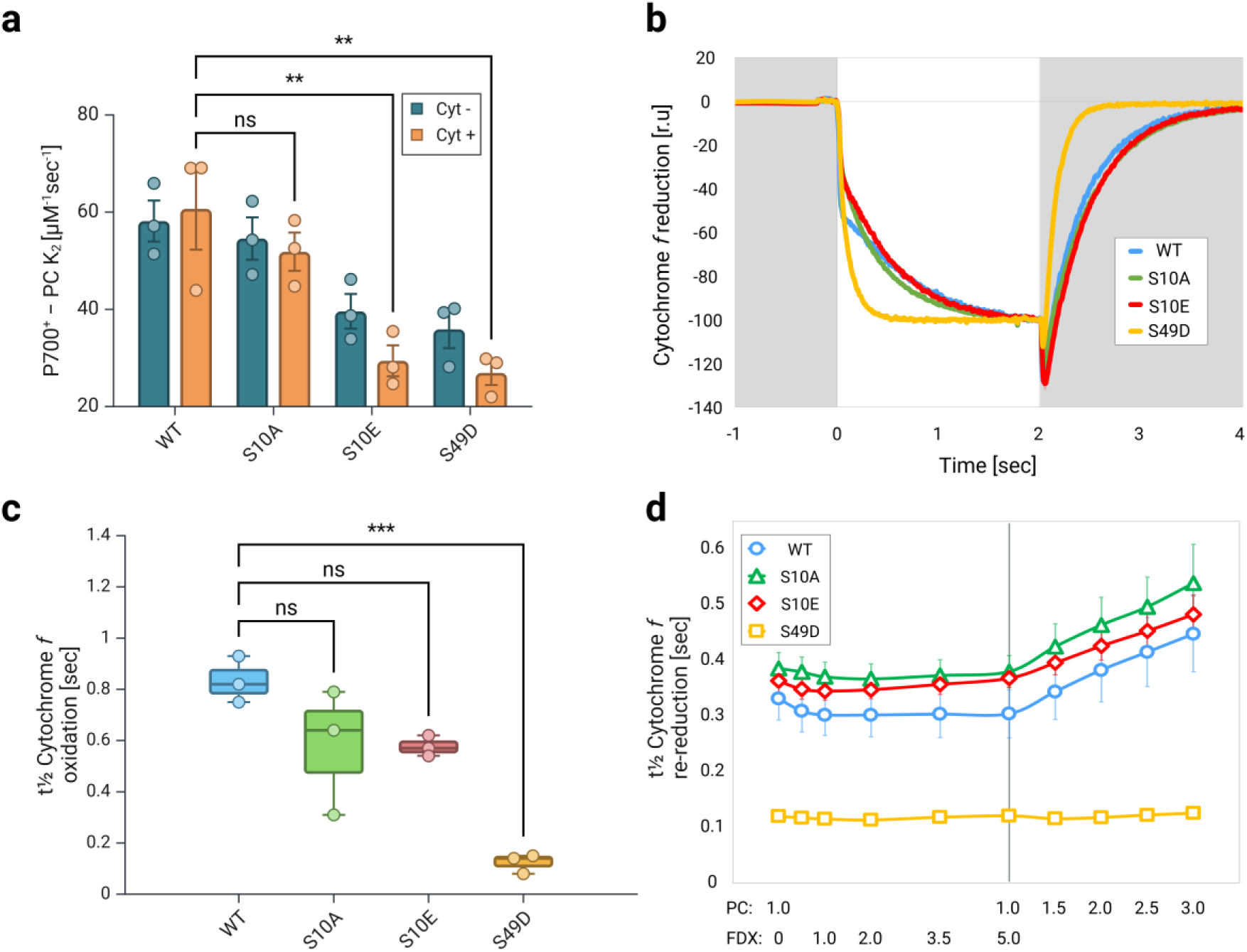
Phosphomimetic PC alters the interaction mode with Cyt*b_6_f*. (**a**) P700^+^ re-reduction assay was conducted in the absence (blue) or presence (Orange) of Cyt *b_6_f*, using different PC mutants (WT, S10A, S10E, S49D). Both S10E and S49D PC peptides showed an additional decrease in K₂ values, indicating a decrease of its apparent concentration and suggesting a stronger interaction with Cyt*f*. (**b**) Comparative Cyt*f* reduction assay, where samples were exposed to 2 sec of illumination (white background), before the light was turned off. (**c**) Further inspection of Cyt*f* oxidation kinetics showed a significant faster oxidation when S49D plastocyanin was added. (**d**) Accordingly, the dark re-reduction of Cyt*f* showed an increased rate when S49D plastocyanin was added, but not due to FDX additions, as depicted for the averaged half-time. Statistical analysis was conducted using a One-way ANOVA with Dunnett multiple comparisons test (for 3 biological repetitions). The illustration and statistical analysis were generated using BioRender.com.

## Discussion

In this work we addressed the putative role of PC phosphorylation in photosynthetic electron transfer. In addition, we investigated the molecular binding between PC and Cyt*f* for *C. reinhardtii*. The electrostatic landscape of PC binding and electron transfer with PSI and Cyt*b_6_f* exhibits striking similarity. In both complexes, positively charged lysine residues in PsaF and Cyt*f* interact with negatively charged aspartate and glutamate residues on PC, aiding in stable complex formation and efficient electron transfer (Figures 1). In plant PSI, the complex between PC and PSI has been structurally described and slight differences have been observed for the complexes in pea and *C. reinhardtii*, as expected from amino acid sequence difference of PC and PsaF (Caspy et al., 2020; Caspy et al., 2021; Naschberger et al., 2022). Likely, due to the amino acid sequence difference of PC and Cyt*f* between vascular plants and green algae, such differences in binding of PC to Cyt*f* could be also anticipated. Indeed, the chemical protein cross-linking coupled mass spectrometry revealed these differences between the vascular plant PC-Cyt*f* interactions (Ubbink et al., 1998) and binding of PC to Cyt*f* in *C. reinhardtii*. The cross-linking data and identified cross-linked peptides between PC and Cyt*f* (Figure. 2) suggested two models, the “Side-on” type (Figure 3c) and the “Head-on” type models (Figure 3d). Both models resulted in a core distance of 10.9 Ȧ (PC-Cu/PetA-heme-Fe). The “Side-on” type (Figure 3c) resembles vascular plant PC-Cyt*f* interactions (Ubbink et al., 1998), yet, it differs on the bond formations with PetA:K121, which positions PetA:K65 in-between PC:D54 and E44 and places PetA:K189 in the proximity of S49. This suggests that phosphorylation of PC:S49 may stabilize the interaction of PC with Cyt*f* and promote electron transfer between the two partner via contact between PetA:K189 and the PC:S49 phosphorylation. Indeed, we observed more efficient oxidation of Cyt*f* via the PC:S49D phosphomimic (Figure 6c) compared to other PC variants. Interestingly, the data also revealed that re-reduction of Cyt*f* via FDX was faster with PC:S49D (Figure 6b). Of particular note are the results in Figure 6d, where increasing PC concentrations for WT PC, PC:S10A, and PC:S10E slowed down FDX-dependent re-reduction of Cyt*f*, whereas the rate remained constant for PC:S49D. Based on our interpretation, we propose that upon exposure to light, PC becomes oxidized at PSI. This oxidized PC is then reduced by Cyt*f* which is in turn re-reduced via FDX. Increasing PC concentrations leads to more oxidized PC at PSI. For WT PC, PC:S10A, and PC:S10E, the oxidation of Cyt*f* appears to be the rate-limiting step in this sequence, thereby slowing down the re-reduction of Cyt*f* as oxidized PC concentration increases. However, in the presence of increasing amounts of oxidized PC:S49D, the oxidation of Cyt*f* proceeds more rapidly, resulting in significantly faster Cyt*f* re-reduction (Figure 6d). This indicates that Cyt*f* oxidation is more efficient with PC:S49D compared to WT PC, PC:S10A, and PC:S10E. In conclusion, these findings support the “Side-on” binding model (Figure 3c) of PC-Cyt*f* interaction, where phosphorylated PC:S49 makes contact with PetA:K188.

It has been described that PC:S49 phosphorylation is induced under HL where at the same time the PC amount is diminished (Younas et al., 2023). Thus, under such *in vivo* conditions the more efficient Cyt*f* oxidation via PC:S49 phosphorylation could help to improve the turn-over of PC between PSI and Cyt*b_6_f*. The PC driven electron transfer between PSI and Cyt*b_6_f* is influenced by the rapid release of oxidized PC from PSI (Drepper et al., 1996; Finazzi et al., 2005) and by the diffusion of PC in the thylakoid lumen that is required for long-range electron transfer between PC and Cyt*b_6_f* complex (Haehnel et al., 1989; Höhner et al., 2020). Such changes in binding affinities could also result differential binding of oxidized and reduced PC as evidenced for PSI (Drepper et al., 1996) and thereby optimize turn-over. Our data suggest that electron transfer between PC and PSI is possibly be influenced by protein phosphorylation. Consistent with this, the phosphomimetic variants PC:S10E and PC:S49D exhibit slower electron transfer rates toward PSI compared to the WT PC (Figure 4c), which was enhanced in the presence of Cyt*b₆f* (Figure. 6a). Although this leads to a two-fold reduction in the rate constant for PSI electron transfer, it does not limit the oxidation of Cyt*f* (Figure 6). In conclusion, we propose that phosphorylation at PC:S49 may be advantageous for optimizing Cyt*f* oxidation under conditions where PC levels are regulated. In this scenario, the “Side-on” conformation is likely favored, though the occurrence of the “Head-on” conformation with non-phosphorylated PC cannot be excluded. These two conformations may have different effects on how electron transfer turnover is catalyzed.

In addition to modulating electron transfer between PC-PSI and between PC-Cyt*b_6_f* complexes via phosphorylation, we cannot rule out the possibility that PC:S10 and/or PC:S49 phosphorylation might also affect its diffusion within the thylakoid lumen, which is essential for long-range electron transfer between PC and the Cyt*b_6_f* complex (Haehnel et al., 1989; Höhner et al., 2020). Mobility shifts in response to phosphorylation are a known process for thylakoid membrane proteins. It is for example well described, that the mobility of LHCII to migrate from grana to stroma membranes increases upon phosphorylation (Allen, 1992) and that this phosphorylation is required for state transitions in green algae and land plants (Depege et al., 2003; Bellafiore et al., 2005). It is possible that phosphorylation, as shown for LHCII, also facilitates the long-range electron transfer from grana localized Cyt*b_6_f* complexes to stromal PSI (Vallon et al., 1991). For Cyt*b_6_f* complexes, the causal relationship between phosphorylation and migration has not yet been demonstrated; however, subunits of the Cyt*b_6_f* complex are also known to undergo phosphorylation (Bergner et al., 2015; Younas et al., 2023). Currently, it is unknown in which sub-compartment PC gets phosphorylated, as no lumen localized protein kinase is known. However, phosphorylation of luminal proteins such as ULP1, CSS1, PSBO and PSBR indicate that phosphorylation is a common phenomenon for luminal proteins in *C. reinhardtii* as well as for *Arabidopsis thaliana* (Spetea and Lundin, 2012; Wang et al., 2014; Bergner et al., 2015; Younas et al., 2023). Therefore, the phosphorylation of PC is not unexpected, although the specific kinase responsible remains unknown. On the other hand, PC and other nuclear-encoded luminal proteins may acquire their phosphorylation during the import process required for translocation from the cytosol into the thylakoid lumen of chloroplasts, as demonstrated for transit peptides in Chlamydomonas (Su et al., 2001). In the light of PC:S49 phosphorylation and its fitting into the binding network via interaction with PetA:K188, it is possible that this mode of binding evolved to accommodate and facilitate fast Cyt*f* oxidation to maintain a sufficiently fast electron transfer between Cyt*b_6_f* complex and PSI. Cyt*f* also underpins the importance of green algal PetA:K121 for rearranging the side-on orientation by forming bonds with PC:D60, and thereby enabling a contact between the phosphorylated PC:S49 and PetA:K188. Notably, in vascular plants, K121 is replaced by glutamine, where it is structurally irrelevant. This is another example of how the electrostatic landscape for PC binding has been optimized to meet the system’s functional demands.

In conclusion, our cross-linking data suggest that in green algae the interaction between Cyt*f* and PC can be formed via two types of orientation, in a phosphorylation-dependent manner. Moreover, we predict a new Chlamydomonas-specific “side-on” model (Figure 3c), in which PC binds to Cyt*f* in a manner distinct from the binding observed in vascular plants (Ubbink et al., 1998).

## Materials and Methods

### PSI Complex preparation

Photosystem I complexes were isolated from *C. reinhardtii*, cc124 WT cells. according to (Iwai et al., 2010). The cell wall was disrupted by nebulizing a dense culture twice in H1 buffer, containing 330 mM sucrose, 5 mM MgCl_2_ solution, pH 7.8 (HEPES 25 mM). Cell extracts were then centrifuged and resuspended in H2 buffer, containing 330 mM sucrose, 10 mM EDTA solution, pH 7.8 (HEPES 5 mM), and pottered, before loaded on a step-sucrose gradient, containing 10 mM EDTA solution, pH 7.8 (HEPES 5 mM), in increasing sucrose concentrations (1.8, 1.3 and 0.5 M). Centrifugation was conducted for 1 hour, in swing buckets (SW32 Ti) at 100,000 g, 4 °C. Thylakoid membranes were then collected from the 1.3-0.5 M sucrose interface and resuspended in a sugar-free buffer, containing 2 mM CaCl_2_. Isolated membranes were then diluted to a chlorophyll concentration of 0.8 mg Chl ml^−1^ and solubilized with 1 % (w/v) n-Dodecyl β-maltoside (β-DDM) by incubating for 20 min on ice. Photosynthetic complexes were separated using a linear sucrose density gradient (average density of 750 mM sucrose, 0.025 % β-DDM). Centrifugation was conducted for 16 hours, in swing buckets (SW41 Ti) at 200,000 g, 4 °C. The gradients were then fractioned, and PSI complexes were concentrated using ultra-filtration columns with a size exclusion of 100,000 MW.

### Cytochrome b₆f Complex preparation

Cyt*b_6_f* complexes were isolated from *C. reinhardtii* strains which express a His-tagged PetA back-crossed to cc125 strains. Thylakoids were extracted as mentioned above, omitting the step-sucrose gradient to improve yields. Solubilization was conducted on crude thylakoid extracts (chlorophyll concentration was set to 2-3 mg Chl ml^−1^) using 0.5 % (w/v) 4-trans-(4-trans-Propylcyclohexyl)-cyclohexyl-α-maltoside (tPCCαM) by incubating for 45 min on ice. The solution was than loaded on a NiNTA column and washed with increasing imidazole concentration (3-18 mM) in 250 mM KCl, 0.01 % tPCCαM, pH 7.8 (HEPES 25 mM), before the complexes were eluted using 300 mM imidazole. Cyt*b_6_f* complexes were then separated using a linear sucrose density gradient (average density of 650 mM sucrose, 0.01 % tPCCαM). Centrifugation was conducted for 16 hours, in swing buckets (SW41 Ti) at 200,000 g, 4°C. The gradients were then fractioned and Cyt*b_6_f* complexes were concentrated using ultra-filtration columns with a size exclusion of 100,000 MW.

### PC mutation and isolation

The heterologous expression of PC was done as described by Hulsker and co-workers (Hulsker et al., 2007). The construction of the WT Pet17b-PETE plasmid was described in (Kuhlgert et al., 2012). Site-directed mutagenesis for the different PC variants was based on (Lei Zheng et al., 2004). Briefly, WT Pet17b-PETE was amplified via PCR using NEB Q5 High-Fidelity 2X Master Mix in 25 µL reaction volumes containing 10 ng DNA template and 0.5 µM of specific primers pairs carrying the new codon for each PC variant (S-to-A mutation: GCC; S-to-E: GAA; S-to-D: GAT); the PCR protocol was set according to the manufacturer’s instructions (amplification was done for 25 cycles with 100 s elongation time each) and annealing temperatures were calculated according to the online NEB Tm calculator. After amplification, 1 µL of PCR product was incubated at RT for 10 min with NEB 10X KLD Enzyme Mix in a 10 µL reaction volume before being used to transform NEB 5-alpha competent *E. coli* cells, for plasmid isolation and Sanger sequencing (Eurofins Genomics). Plasmids with the correct sequence were used to transform NEB BL21(DE3) competent *E. coli* cells for recombinant protein production. Following O.N expression (LB medium, 1mM IPTG, 100 µgr/mL Amp, 0.1 mM CuSO₄), the cells were harvested and resuspended in lysis buffer (KCl 10 mM, Cu(NO₃)₂ 10mM, PMSF 1mM and tricine 10 mM pH 7.8). The cells were then sonicated for a total of 180 sec (Branson 250 Digital Sonifier w/ Prob, Marshall Scientific), supernatant was separated via centrifugation (20 kG, 30 min) and loaded on an anion exchange column (DEAE Sepharose CL-6B, GE Healthcare). The columns were washed with KCl 50 mM and eluted with 400 mM KCl. The proteins were then concentrated, and salt concentration diluted back to 10 mM, before loading them on a size exclusion chromatography (Superdex 65 10/300 GL on an ÄKTA pure system). The concentration of PC was determined spectroscopically (at 597 nm with an extinction coefficient of 4.7 mM^−1^ cm^−1^ (Yoshizaki et al., 1981)).

### FDX isolation

FDX (plasmid: Pet21b:FDX:TEVc:MBP:His6) and Tobacco Etch Virus (TEV) protease (plasmid: pMHTDelta238) heterologously expressed in *E. coli* were a kind gift of Prof. Dr. Iftach Yacobi from Tel-Aviv University. Expression and purification was conducted as described in (Marco et al., 2018).

### Cross-linking

Cyt*b_6_f*-PC Cross-linking was performed as described in (Naschberger et al., 2022). Isolated PC (100 µM in 40 µL) was pre-activated with a 5 mM 1-ethyl-3-[3-dimethylaminopropyl]carbodiimide (EDC) and 10 mM sulfo-N-hydroxysuccinimide (NHS) solution, pH 6.5 (MOPS 10 mM) for 20 min at room temperature. 5 mM ferricyanide was then added before the cross-linker was removed and the buffer was exchanged to Ho buffer, pH 7.5 (HEPES 30 mM) via a PD G25 desalting column followed by ultrafiltration with a 0.5 mL centricon (regenerated cellulose: 3,000 MWCO). Isolated Cyt*b_6_f* complexes (2.7 µM in 100 µL) were pre-reduced using 5 mM ascorbate, in Ht buffer containing 0.01 % tPCCαM, pH 7.5 (HEPES 30 mM). Ascorbate was then removed using a PD G25 desalting column followed by ultrafiltration with a 0.5 mL centricon (regenerated cellulose: 100,000 MWCO) and set back to 100 µL in Ht. Activated PC was then added, in addition to 60 µL of Hm buffer containing 10 mM MgCl_2_ and 0.0167 % tPCCαM (to adjust final detergent concentration), pH 7.5 (HEPES 30 mM). The mixture was finally cross-linked for 45 min at room temperature, before samples were taken for further analysis.

### SDS-PAGE

For SDS–PAGE 30 μL of the cross-linked mixture (approximately 42 pmol of total protein) were sampled, supplemented with loading buffer (Glycerol 8%, SDS 1.33%, Servablue G 0.007%, Bromphenolblue 0.007%, Tris 35 mM pH 8.0), sodium dithionate 167 mM, and sodium carbonate 167 mM, and incubated at 65 °C for 20 min. Proteins were separated on 13% (w/v) SDS–PAGE, with a stacking gel of 4% (w/v) (Laemmli, 1970). Gels were either stained with Coomassie Brilliant Blue (R-250) or blotted onto nitrocellulose membranes (Amersham). Autofluorescence detection of heme-containing proteins was done in the absence of antibodies, using a Supersignal West Pico Plus chemiluminescent substrate (Thermo Scientific) for 10 min. The proteins were also then incubated with Cyt*f* antibodies (Agrisera, Anti-Cyt f, AS06 119), which resulted in the same signal bands.

### Mass Spectrometry preparation and analysis

Bands from relevant Coomassie-stained gels were cut, de-stained and typically digested according to (Shevchenko et al., 2006). Cross-linked peptide samples were analyzed using an LC-MS/MS system comprising an Ultimate 3000 nano HPLC (Thermo Fisher Scientific, Waltham, MA, USA) connected via an ESI interface (Nanospray Flex, Thermo Fisher Scientific) to a Q Exactive Plus mass spectrometer (Thermo Fisher Scientific). Samples were reconstituted in solvent A1 (0.05% trifluoroacetic acid (TFA)/2% acetonitrile (AcN)/ultrapure water) and loaded onto a trap column (C18 PepMap 100, 300 µM × 5 mm, 3 µm particle size, 100 Å pore size; Thermo Fisher Scientific) at a flow rate of 10 µL/min for 3 minutes using solvent A1. Peptides were then eluted in backflush mode from the trap column to the separation column (PepSep C18, 75 µm × 15 cm, 1.9 µM particle size, Bruker) at a flow rate of 250 nL/min. Eluents for peptide separation included 0.1% formic acid in ultrapure water (A2) and 80% AcN/0.1% formic acid in ultrapure water (B). The elution gradient was as follows: 2.5% to 40% B over 50 minutes, then 40% to 99% B over 5 minutes, with 99% B maintained for 10 minutes. MS full scans (m/z 350–1450) were recorded at a resolution of 70,000 (FWHM at 200 m/z. Fragmentation spectra (MS2) were acquired at a resolution of 17,500 (FWHM at 200 m/z) using a data-dependent method, where the 12 most intense ions from each full scan were fragmented via higher-energy c-trap dissociation (HCD) at a normalized collision energy of 28 and an isolation window of 1.5 m/z. The automatic gain control (AGC) targets were set to 3e6 for MS full scans (MS1) and 5e4 for MS2, with an MS2 intensity threshold of 8.3E3. Maximum injection times were 50 ms for MS1 and 60 ms for MS2. Ions with unassigned charge states or those with charge states of 1, 2, or ≥8 were excluded from fragmentation.

### Analysis of PC mutants

Instruments and buffers were the same as described above. Trap column loading was performed at 25 µL/min for 1.5 min. Peptide separation was carried out using a PepSep C18 column (75 µm × 50 cm, 1.9 µM particle size, Bruker) operated at a flow rate of 250 nL/min. The elution gradient was programmed as follows: 2.5% to 5% B over 5 minutes, 5% to 35% B over 105 minutes, 35% to 99% over 18.5 min, followed by a hold at 99% B for 20 minutes. MS full scans (m/z 400–1800) and fragmentation spectra (MS2) were acquired at a resolution of 70,000 (FWHM at 200 m/z) and 35,000 (FWHM at 200 m/z), respectively. Peptide fragmentation was conducted using a data-dependent approach, fragmenting the 12 most intense ions from each full scan through higher-energy c-trap dissociation (HCD) at a normalized collision energy of 28 and an isolation window of 1.5 m/z. Automatic gain control (AGC) targets were set at 3e6 for MS full scans (MS1) and 1e5 for MS2, with an intensity threshold of 1E4 for MS2. Maximum injection times were 50 ms for MS1 and 120 ms for MS2. Ions with unassigned charge states, or those with charge states of 1, 2, or ≥8, were excluded from fragmentation.

### Cross-link identification

MS raw files were analyzed using MaxQuant 2.4.14.0 (Cox and Mann, 2008) against the polypeptide sequences of *Chlamydomonas reinhardtii* proteins forming the Cyt*b_6_f* complex (PETx). When PC mutants were analysed, corresponding mutated sequences were used in addition. Default search settings for identification of peptides cross-linked using EDC were applied. Minimum peptide length was 6. Carbamidomethylation of cysteine was set as fixed modification. Variable modifications included N-terminal acetylation and methionine oxidation. Cross-linked peptides were filtered to maintain a false discovery rate (FDR) of 0.01. The mass spectrometry proteomics data have been deposited to the ProteomeXchange Consortium via the PRIDE partner repository with the dataset identifiers with accession nr-PXD060078. A detailed list of all identifications of cross-linked peptides is also available as Supplemental File 2.

### Sequence alignment

Sequence alignments were conducted by taking all of the available sequences from Uniprot.org (https://www.uniprot.org/), using the keywords: PETE (PC), PsaF, and PetA. The data was filtered for duplications of identical sequences (in cases where the same organism has multiple annotations). Organisms were clustered to families and aligned according to conserved regions, with highest similarity. Reported sequence is numbered according to the sequences of *C. reinhardtii* (PC-ID: P18068 starting from D48 = D1 here, PsaF-ID: P12356, starting from D63 = D1 here and PetA-ID: P23577 starting from Y32 = Y1 here). Moreover, the most frequent amino acid is presented (for a full alignment see, Supplemental File 1), when alignments are based on structural localization of the residues.

### Structural modelling

Docking of PC and PetA was performed using the ClusPro.2 docking server (https://cluspro.bu.edu) (Xia et al., 2016; Kozakov et al., 2017; Desta et al., 2020) . The structure of Cyt*b_6_f* (PDB: 1Q90, (Stroebel et al., 2003)) was used as a receptor and the structure PC (PDB: 7ZQE, (Naschberger et al., 2022)) was used as a ligand. Restrains were determined using provided plug-in (https://cluspro.bu.edu/generate_restraints.html), optimal dockings were filtered base on core distances, between the Cu ligand of PC and the Fe ligand in the center of Heme *f*. For a full list of docking attempts and detailed restraints please see Supplemental File 3.

### Fast optical spectroscopy measurements

Isolated PSI complexes from *C. reinhardtii*, WT strain cc124. were mixed in the presence of PC (0.3-6.0 µM) and 331 nM PSI, solubilized in 7.5 mM KCl, 2.5 mM MgCl_2_, 10mM ascorbate, 1 mM methyl viologen and 0.5 mM DAD, pH 7.0 (MOPS 5mM), when mentioned, isolated Cyt*b_6_f* complexes were added (2.1 µM). Absorbance was measured post a Laser flash using ‘Joliot type Spectrophotometer’ (JTS -150, Biologics) supplied with a ‘Smart-Lamp’ with a dual measuring light usage (705-740 nm) and adequate detector filters (P700). To conduct measurements of Cyt*f* fast kinetics, we used a set-up of triple measuring light (where 554 nm indicates on Cyt*f* absorbance with a base-line drawn between 546 and 573 nm), and adequate detector filters (BG-39). Actinic light was supplied by an orange ring (630 nm, 300 µE m^−1^ s^−1^). Each test was composed of 4 technical repetitions which were averaged. Each type of PC was tested using three different PSI complexes, from individual isolations. For an elaborated data set and analysis, see Supplemental File 5.

## Acknowledgements

This research was financed by the Deutsche Forschungsgemeinschaft (DFG HI739/13-1/2, DFG / 9-1/-2 and 25-1 in DFG FOR 5573/1), in association with the University of Okayama, Japan. Y.M. acknowledges the Alexander von Humboldt Foundation for a Research Fellowship for postdocs (1219125).

## Author Contributions

Research design was conducted by Y.M., S.K., A.V.M. and M.H. Complex and protein construction and purifications were conducted by Y.M., D.W., S.K., M.Y. and A.V.M., Cross-linking experiments and sample preparation were conducted by Y.M. and D.W. Mass-spectrometry studies and analysis were conducted by M.S. Complex interaction modeling was conducted by Y.M. Kinetic studies were conducted by Y.M. and S.K. the manuscript was written by Y.M. and M.H.

## Data Availability

All data will be available online once accepted. Upon submission, the authors attached additional files which include all raw data and analysis. For any additional request, please contact the corresponding author.

