## Supplemental Figure 1 for "New Insights into Plastocyanin– Cytochrome *b_6_f* Formation: the Role of Plastocyanin Phosphorylation": Supplemental Figure 1- Models zoom on residues with legend.pdf

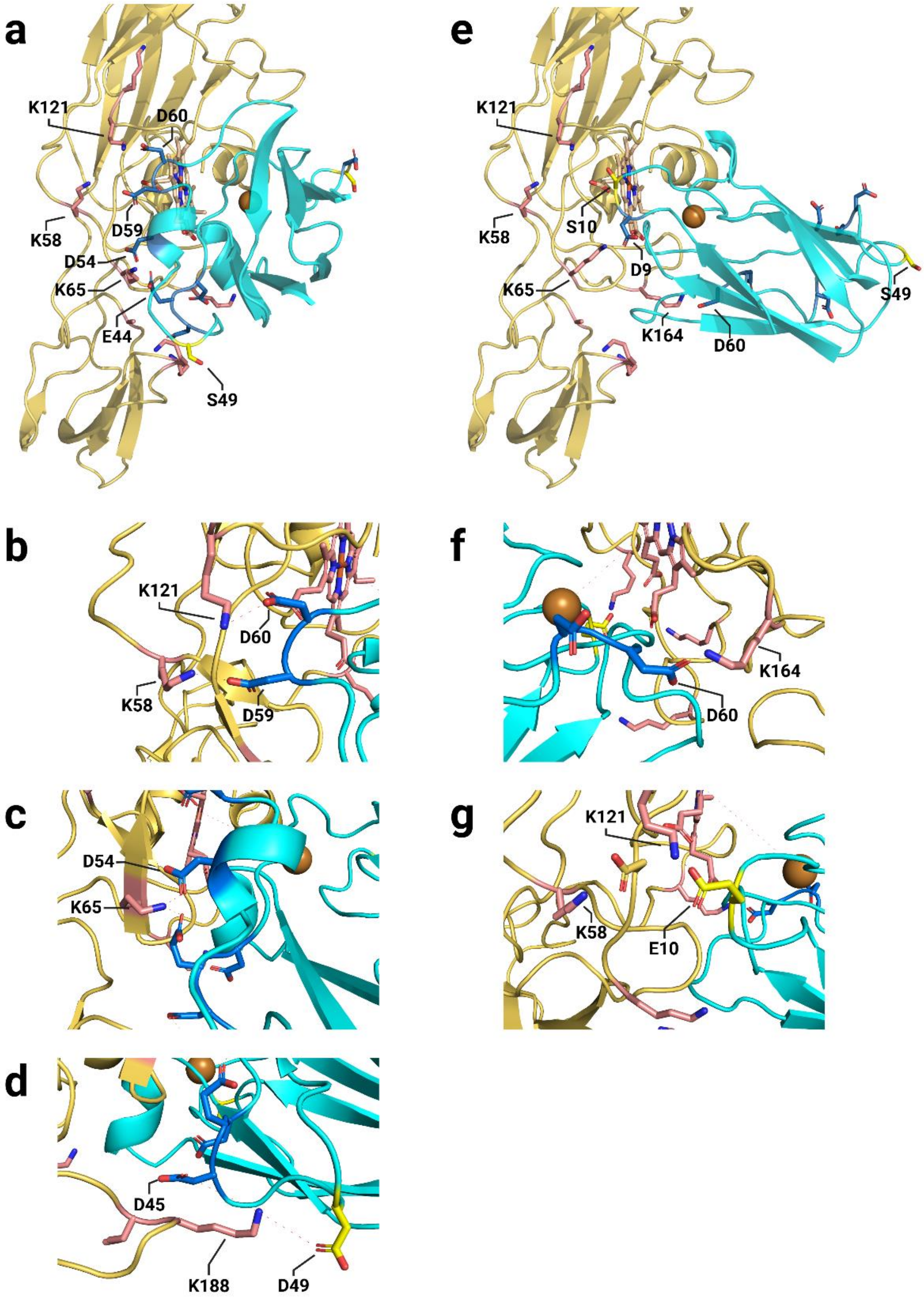


**Supplemental Figure 1. Structural modelling of Cyt*f* and PC in complex.** Illudstration of the two models as presented in Figure 3. (**a**) This model takes into consideration the fact that algal PetA has an additional Lys at position 121, and that it was cross-linked to PC:D59/D60 (**b**). Additional interactions are observed between PC:D54 and PetA-K58/K65, PC:D43/E44 and PetA-K188/K65 (**c**). In addition, PC:S(D)49 seems to be in the proximity of PetA-K188 (**d**) and given a phosphorylation form might increase the stability of the complex formation. Core Cu-Fe distances are predicted to be 10.9 Å. (**e**) the second model takes into consideration a possible interaction between PC:D60 and PetA-K164 (**f**). This model also predicts an interaction between PetA-K121 and PC-S(E)10. All models are available on Supplemental File 4. The illustration was generated using BioRender.com.
